# Active dendrites enable robust spiking computations despite timing jitter

**DOI:** 10.1101/2023.03.22.533815

**Authors:** Thomas SJ Burger, Michael E Rule, Timothy O’Leary

## Abstract

Dendritic action potentials exhibit long plateaus of many tens of milliseconds, outliving axonal spikes by an order of magnitude. The computational role of these slow events seems at odds with the need to rapidly integrate and relay information throughout large nervous systems. We propose that the timescale of dendritic potentials allows for reliable integration of asynchronous inputs. We develop a physiologically grounded model in which the extended duration of dendritic spikes equips each dendrite with a resettable memory of incoming signals. This provides a tractable model for capturing dendritic nonlinearities observed in experiments and in more complex, detailed models. Using this model, we show that long-lived, nonlinear dendritic plateau potentials allow neurons to spike reliably when confronted with asynchronous input spikes. We demonstrate this model supports non-trivial computations in a network solving an association/discrimination task using sparse spiking that is subject to timing jitter. This demonstrates a computational role for the specific time-course of dendritic potentials in situations where decisions occur quickly, reliably, and with a low number of spikes. Our results provide empirically testable hypotheses for the role of dendritic action potentials in cortical function as well as a potential bio-inspired means of realising neuromorphic spiking computations in analog hardware.

## Introduction

Across species, many types of neurons possess active dendrites that produce nonlinear responses to synaptic input [38, 50, 25, 10]. The computational role of these nonlinearities is diverse and likely impacts the function of the wider neural circuit they inhabit. Some of the most intensely studied examples of dendritic excitability are found in cortical excitatory neurons, which produce regenerative dendritic potentials – ‘dendritic spikes’ – in response to excitatory synaptic drive [50, 63, 46].

Dendritic spikes in excitatory cortical neurons last for many tens of milliseconds [63, 62, 61, 51, 47, 46]. This timescale is conspicuous, being many times longer than the fastest signalling processes in the nervous systems, including Excitatory Post-Synaptic Potentials (EPSPs) and axonal spikes (Figure 1). Reconciling the relatively slow time course of dendritic potentials with rapid signalling and computation therefore poses a challenge, particularly when such computations may involve relaying information over multiple brain areas [70]. Furthermore, the duration of dendritic events incurs heavy energetic costs, because dendritic currents contribute significantly to the ATP budget of the brain [5]. What computational benefit might justify these signalling and metabolic costs?

**Figure 1:**
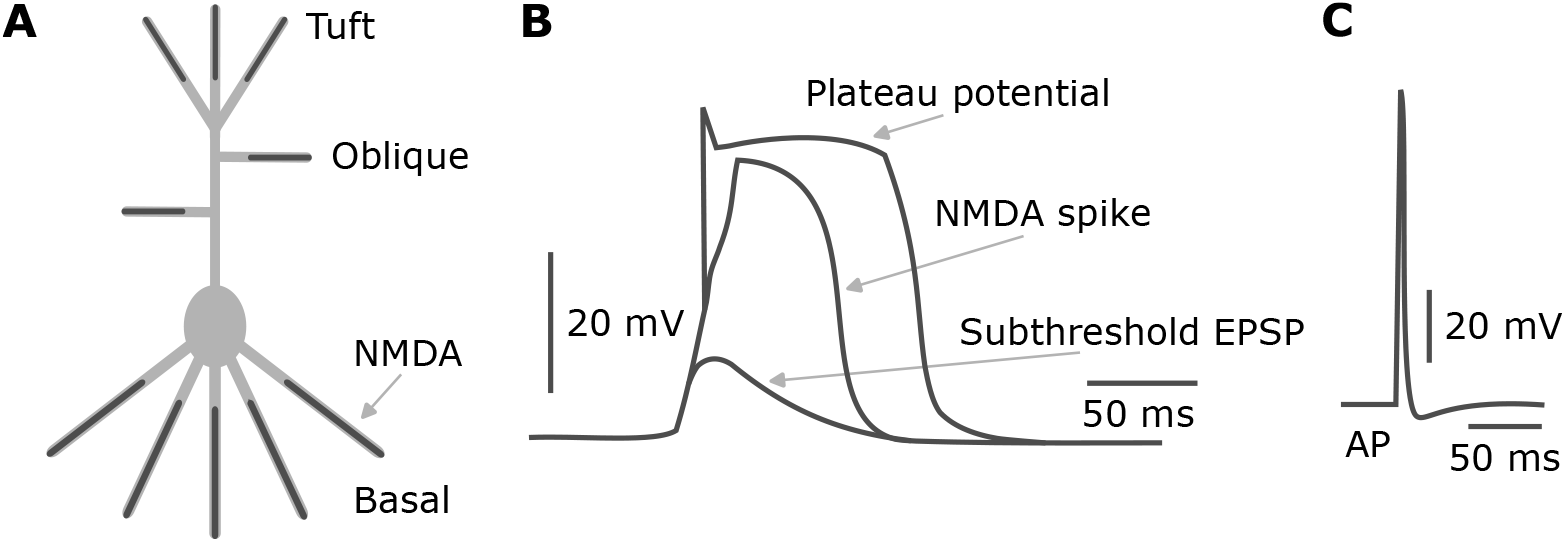
Reproduced from [3]. Dendritic NMDA-dependent action currents. **(A)** Long lived voltage transients can be initiated by NMDA receptors located within distal dendritic branches. **(B)** If the input to a dendritic branch with NMDA receptors is sufficiently strong, it can cross a threshold to produce an NMDA spike (middle trace). The NMDA response to inputs is super-linear, until it saturates in a plateau potential (top trace). **(C)** A somatic spike mediated by voltage-gated sodium channels. Note the order of magnitude difference in timescales with (b).

We propose that the duration and threshold-like properties of dendritic currents support robust computation in the face of spike timing jitter by holding a trace of recent inputs. This is especially relevant to integration of inputs during high conductance states that are prevalent in vivo, where a typical neuron receives significant synaptic drive. In these states, the effective membrane time constant can be extremely short, and varies substantially depending on synaptic input [16, 42, 57]. As a consequence, computations that rely on passive summation of multiple inputs would place strong constraints on spike timing precision. Dendritic action potentials, by contrast, have a consistently long durations that are ensured by the gating properties of voltage gated ion channels and NMDA receptors [62, 55, 12, 3]. These properties are largely determined by the amino acid sequence of receptor and channel proteins that are expressed in dendrites [52, 51, 47]. Dendritic properties thus appear to be specifically tuned at the molecular and genetic level to produce localized, suprathreshold events that outlive rapid membrane fluctuations.

Numerous studies point out that nonlinear summation in dendrites can make neurons computationally equivalent to entire networks of simplified point models, or ‘units’ in a traditional neural network [11, 26, 7, 46, 47, 52, 56, 58, 59]. Other work has shown that the dynamic properties of dendritic action potentials enrich computational and signal processing capacity by providing additional long timescales over which input-driven membrane potential or synapse dynamics evolve [27, 59, 9, 45, 58, 30, 4, 20]. These ideas and the specific examples that support them are complementary to what we are proposing here. With the dendritic potential as a backbone, our work adds to the computational repertoire by allowing neurons to achieve robust computation even when there is high variability in the arrival times of inputs.

We extract core features of the biophysics of dendritic integration to construct and analyze a simplified model. With this model we demonstrate that rapid computation remains possible – and is in fact facilitated – by dendritic transients that exceed the integration time constant of single neurons. This idea is not in conflict with other hypothesized functions of dendritic transients, including the induction of synaptic plasticity [39, 29]. The extent of interaction between plasticity and the mechanism we propose here is thus an important question for future work. We focus on computations that take place on the most rapid timescale, because short integration windows are necessarily more sensitive to timing jitter.

A side product of our analysis is the interpretation of rapid cortical computations as being binary in nature, with each neuron participating with at most one spike per computation. A number of studies find empirical evidence for such an operating regime in different parts of the nervous system, such as sensory decision making in complex visual and auditory scenes [17, 32, 69, 70]. Within this operating regime, and possibly more generally, we show how dendritic potentials allow non-trivial, robust and rapid spiking computations at a network level. Our work therefore suggests that long-lived dendritic potentials can paradoxically assist in the most rapid computations possible in a spiking network.

## 1 Results

### 1.1 Abstract model

Key features of NMDA action currents are their long duration and their super-linear integration of inputs [50]. Figure 2A and B show a recording of an NMDA spike from a cortical neuron in a rat, from [23]. [23] triggered an NMDA spike by glutamate micro iontophoresis while recording at the soma and the indicated (red, blue) sites in the dendrite. The voltage response (Figure 2B) reveals the orders-of-magnitude difference in timescale of an NMDA spike (left) compared to a sodium action potential in the soma (right). We took their extensive biophysical computational model (85 compartments, 439 segments—for details see [23] and subsection 3.1) and simulated glutamate releases 50 ms apart in the three dendritic sites indicated in Figure 2C, thereby triggering three NMDA spikes at those sites, analogous to the experiment of [54]. Despite these dendritic spikes being initiated at different times, they still sum in the soma, leading to a sodium action potential there (Figure 2D).

**Figure 2:**
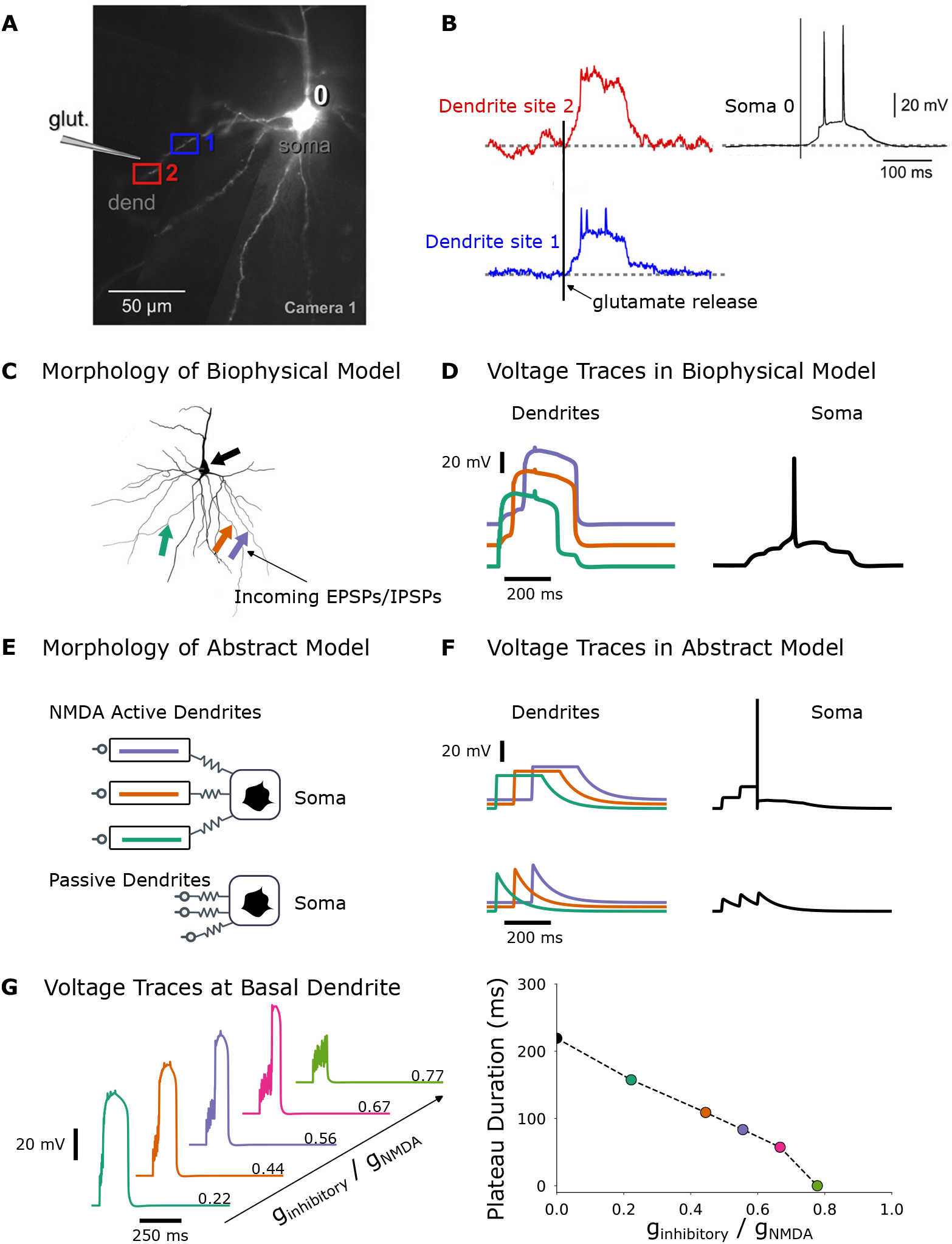
**(A)** Reproduced from [23]. Fluorescent dye fill image of a layer V pyramidal neuron recorded intracellularly in a rat cortical slice [23]. Two dendrites are marked where an NMDA spike is triggered through release of glutamate. **(B)** Voltage traces of the NMDA spikes of the dendritic patches marked in A. Note the long duration of the NMDA spike compared to the somatic sodium spikes evident in the top-left panel. **(C)** The morphology of the biophysical model used for simulating detailed NMDA plateau potentials. The arrows mark the positions where glutamate release was simulated. **(D)** The three traces of the NMDA spikes triggered at the sites marked in C, and the resulting somatic spike. **(E)** The morphology of the abstract model, with and without active NMDA dendrites. **(F)** The voltage traces of the abstract model, with and without plateaus. Because of the extended time duration of the plateau potentials, they are summed to reach threshold. In the case where the plateau potentials are absent, they do not sum due to the short membrane time-constant of the soma. **(G)** Voltage traces of a basal dendrite with an NMDA spike, in the biophysical model, with an increasingly strong inhibitory current added (left). The plateau duration decreases linearly for a linear increase in the inhibitory conductance (right).

We developed an abstract model of these NMDA spikes to capture the essence of their role in circuit computations and for computational expediency. The model consists of a somatic compartment coupled passively to multiple dendritic compartments, each of which corresponds to a single branch on the dendritic tree (Figure 2E; for details see subsection 3.2). We model NMDA spikes by thresholding the voltage of a leaky dendritic compartment. When the dendritic voltage exceeds threshold it remains depolarized for some time before returning to rest (Figure 2F). The voltage dynamics are thus parametrised by the threshold and duration of the NMDA spike. We refer to this behavior as “Leaky Integrate-and-Hold” (LIH). It captures the salient features of the NMDA spikes, namely the threshold plus saturation of the super-linear integration, and the long-lived plateau of the dendritic voltage.

Due to passive coupling between compartments in the model, excitation in the dendrites depolarises the soma membrane potential, potentially leading to “axonal” output spikes. We do not model an axonal compartment; instead we implement standard leaky integrate-and-fire (LIF) dynamics in the somatic compartment. A detailed description of the model, along with links to code, is provided in the methods.

We compared the behaviour of our simplified model with that of the full, detailed biophysical model of Figure 2C,D. The plateau potentials in the abstract model have a qualitatively similar effect on somatic membrane potential as the NMDA spikes in the biophysical model: Figure 2F, top, shows that spikes arriving at different times are summed in an integrate and hold-like manner.

We compared this to a situation where all inputs arrive at a soma with standard LIF dynamics and a 10 ms membrane time constant. This time constant is consistent with the high-conductance state of pyramidal neurons in the cortex [8]: inputs decay after 2–3 ms, and fail to sum to spike threshold (Figure 2F, lower).

To partially account for effects of inhibition, we assessed the robustness of dendritic plateaus to tonic inhibitory conductance. As can be seen in Figure 2G, dendritic plateaus survive inhibitory conductance up to values where the total conductance is roughly equal (see also [18]). Thus, to a crude approximation, dendritic potentials provide an integrate-and-hold mechanism that could function e.g. in the balanced regime observed in cortical circuits. In the present study we did not attempt to account for temporal variation in inhibition, which will likely play a role in providing further spike synchrony, among other things. This is an important issue for future work. In the scope of what remains here we want to ask if Leaky-Integrate-and-Hold (LIH) can easily and plausibly facilitate network computations with spikes.

### 1.2 Single Neurons Struggle to Integrate Asynchronous Spikes

The simplified model captures a key feature of the detailed biophysics of pyramidal neuron dendrites: the ability to integrate and hold inputs for a duration exceeding the membrane time constant. We hypothesized that this feature would be useful in situations where neurons need to integrate asynchronous input and reliably threshold it despite fluctuations in arrival times of the input.

In effect, each dendrite is performing a binary classification on its inputs. If input spikes arrive in a narrow time window, reliably integrating them is trivial (Figure 3A,B left). However, millisecond-scale synchrony is unlikely in a large network that is subject to uncertainty and noise. Empirically, spike timing jitter is commonly observed at the population level.

**Figure 3:**
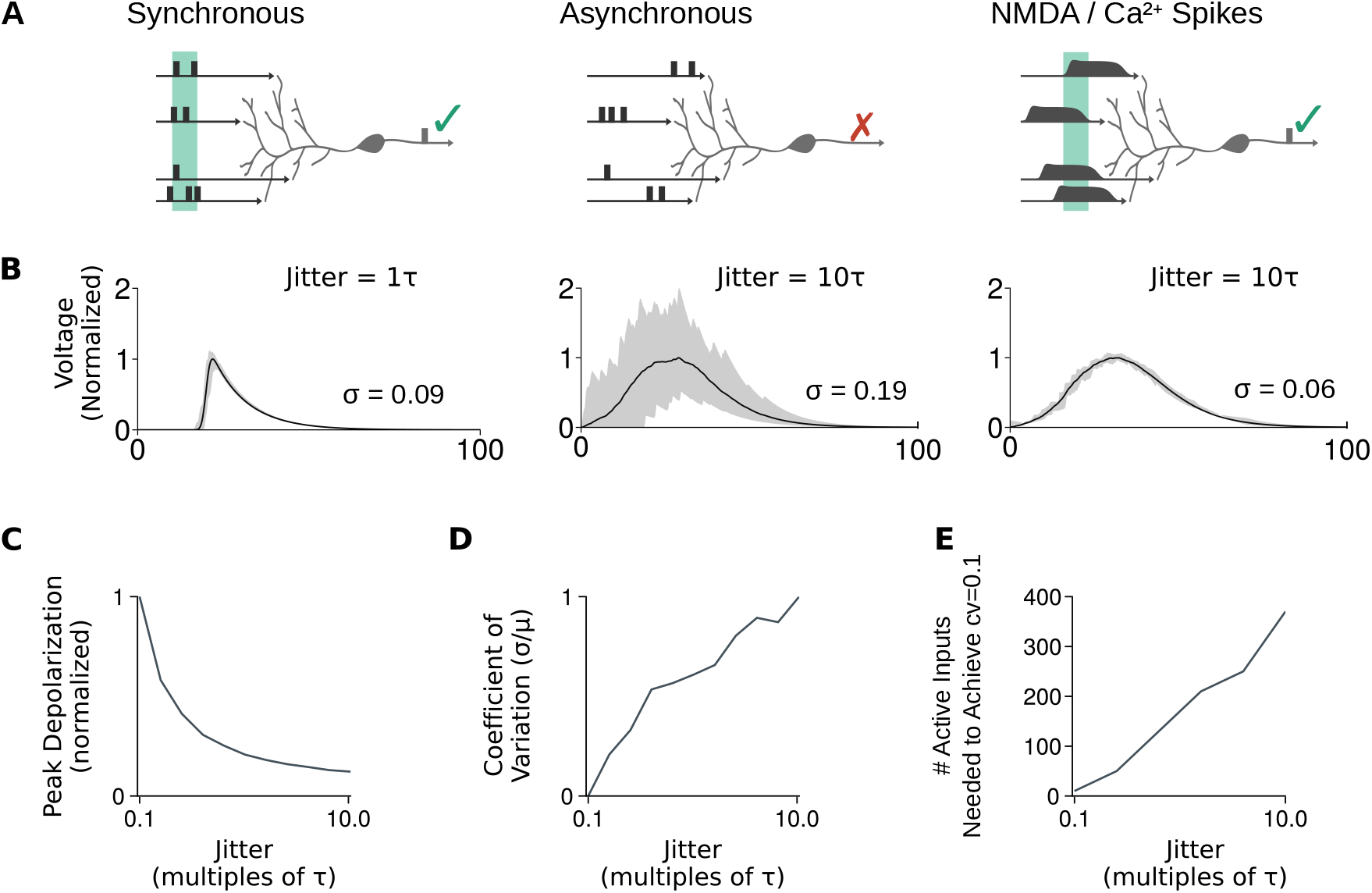
**(A)** Neurons integrate inputs and compare the result to a firing threshold, which is comparable to performing a binary classification. When inputs are synchronous this can be done with a low number of spikes (left). But when spike timing is unpredictable (middle), this falls apart. Extended depolarizing potentials within dendritic compartments acts as a hold mechanism, allowing asynchronous events from different compartments to summate (right). **(B)** Simulation of summed voltage for 10 dendritic compartments, for small amounts of input-event timing jitter (on the order of one EPSP duration *τ ∼* 10 ms; left), and larger amounts jitter (10*τ*, middle). Increased jitter increases the variability of the net depolarization. Extended depolarizing potentials on the duration of *∼* 20 ms reduce the variability in net depolarization. Voltage values were normalized by dividing by the maximum of the mean, and the shaded are shows the inner 90% of the resulting distribution of the voltage. **(C)** Increased jitter in the timing of input events reduces the net summed depolarization. **(D)** Increased jitter in the timing of input events increases the variability in membrane voltage depolarization. **(E)** Variability can be controlled by increasing the number of inputs, but this is not cost-effective.

To illustrate the severity of this problem, we modelled a single neuron using our abstract model, and fed it input spikes. We drew the times of these input spikes from a normal distribution, and varied the degree of input synchrony by changing the standard deviation of this normal distribution. We took the standard deviation to be a function of the membrane time constant *τ*, which defines the timescale of the neuron dynamics.

Spikes arriving even slightly out of sync with each other introduce noise in the membrane potential of the receiving neuron (Figure 3B,D), which can lead to the neuron failing to spike when it should have, or vice-versa. Asynchrony reduces the effective drive of the inputs (Figure 3C), which means that a failure to spike will occur more often than an errant spike. This loss of drive could be compensated by lowering the postsynaptic cell’s threshold, but variability due to jitter remains. This is shown in Figure 3D, where we used the coefficient of variation of the peak membrane potential (standard deviation divided by the mean) to summarize the membrane-voltage uncertainty. This grows with increasing input-timing jitter.

Having extended NMDA spikes remedies these issues (Figure 3A). Because of the extension of the time-constant of the spikes, the uncertainty is filtered out, and the spikes are integrated as if they had arrived synchronously.

It is worth noting that these problems can also be addressed by increasing the number of inputs, thereby reducing the uncertainty through averaging. However, many inputs are required to keep the uncertainty in the membrane potential low; Figure 3E shows that there is a linear relationship between the spike timing jitter in the inputs, and the number of input spikes necessary to have low uncertainty. Even for a relatively low amount of jitter in the input spike timings of 10 ms, the number of inputs required is in the hundreds. Furthermore, it is only possible to average-away timing jitter if timing variations are uncorrelated. This need not be the case, especially if timing jitter arises from variable conduction delays from common sources.

### 1.3 Active Dendrites confer Robustness in Spiking Computations

So far we have shown how a mechanism extracted from detailed biophysics naturally extends the integration window for excitatory inputs. While this might be useful in principle, it remains for us to show how such dynamics can permit non-trivial computations in a network.

In our model, the crucial difference between summation at the soma and summation at the dendrite is that each dendrite can sum subthreshold inputs passively and independently, while the soma is summing sustained plateau potentials from all dendrites that happen to be active. We have made fairly simple assumptions that the dendrite has linear properties beneath the threshold for a dendritic spike, and a relatively short time constant. If neither of these assumptions hold, then dendrites might have even more robust integration properties than our model assumes. In this sense our claims and results in this section are conservative.

We assumed that inputs to a network arrive at the dendrites within some time window, and their combined depolarisations are either sufficient to elicit a dendritic spike or not, as shown in Figure 3. We consider *fast computations*, that is, a regime where the window in which spikes arrive is small, but not so small as to be equivalent to assuming perfect synchrony.

In this regime, each dendrite integrates over a time window and either reaches threshold or does not. Because we assume spike timing jitter in all inputs, each dendrite might reach threshold at different times for different presentations of the same nominal input. However, because dendritic spikes are sustained, jitter in the onset of these events across an entire cell has relatively little effect on whether the soma reaches threshold or not. This effect should confer spike timing robustness at the network level, which is the main claim we will test.

Before describing the implementation of the model and the results, we introduce a useful interpretation of the operating regime of the network. Network dynamics on short timescales can be interpreted as a binary computation, where incoming connections can be represented with a 1 (a presynaptic spike arrives) or a 0 (absence of a presynaptic spike), and the dendrite in turn produces a 1 (it fires) or a 0 (it does not). Connections between dendrites and soma are interpreted analogously: the dendrites produce 1s or 0s, and the soma sums these and compares the result to a firing threshold, thereby computing a 1 or 0. Interestingly, neurons and dendrites operating in this regime have been observed empirically, see e.g. [17, 71].

We used this binary interpretation in both a philosophical and practical sense in our spiking model. Philosophically, the binary analogy provides a clean intuition for how fast computations operate, with each unit (i.e. neuron) only having sufficient time to spike once during a single “pass” or “cycle” of a computation. Such situations do appear in biology; see e.g. [69] for an example where neurons at each synaptic stage have about 10 ms to process presynaptic spikes and fire during an animal’s reaction time, leaving room for the receiving and firing of about one action potential per unit.

On a practical level the binary interpretation gives us a means for finding synaptic weights that allow us to train a network to perform a non-trivial computation, then test its robustness to timing jitter. We remind the reader that the focus of our investigation is not on the training or learning procedure, so the fact that we can train binary networks and use the weights in a dynamic, spiking network is not strictly relevant to the biology. However, it may give a powerful practical means for optimising hardware implementations of dynamic spiking networks. It may also hint that biological learning rules can operate in a somewhat equivalent manner.

We now outline the implementation. We built a Spiking Neural Network (SNN) where the individual neurons consist of our abstract neuron model. We constructed a separate Binary Neural Network (BNN) with the same number of equivalent units, and trained it using standard optimisation methods to perform a classification task (see: subsection 3.4 for details). The BNN is a static network where each unit’s state is either 0 or 1. BNNs can thus be regarded as the saturated limit of regular sigmoidal networks, i.e. with weights of large absolute value [49]. As an aside we point out that BNNs are not restrictive computationally: any computable function can be approximated and implemented with a BNN [48, 66].

The task we train the BNN for is shown in Figure 4A. The 2D input points (three classes, Figure 4A) were first projected onto a binary feature space, to obtain three binary vectors; each 13-D binary vector responding to a specific class The dimensionality of 13 was chosen because this was the lowest dimensionality in which the binary network could still cope with the loss of information due to the binarization of the continuous coordinates. If the *i*^th^ input of the binary vector was a 1, a randomly generated timepoint *t*_*i*_ was added to produce an input spike (*i, t*_*i*_), meaning that input neuron *i* was fed an impulse so that it would spike at time *t*_*i*_. If the *i*th element of the binary vector was 0, it meant that neuron *i* would not fire for that input vector.

**Figure 4:**
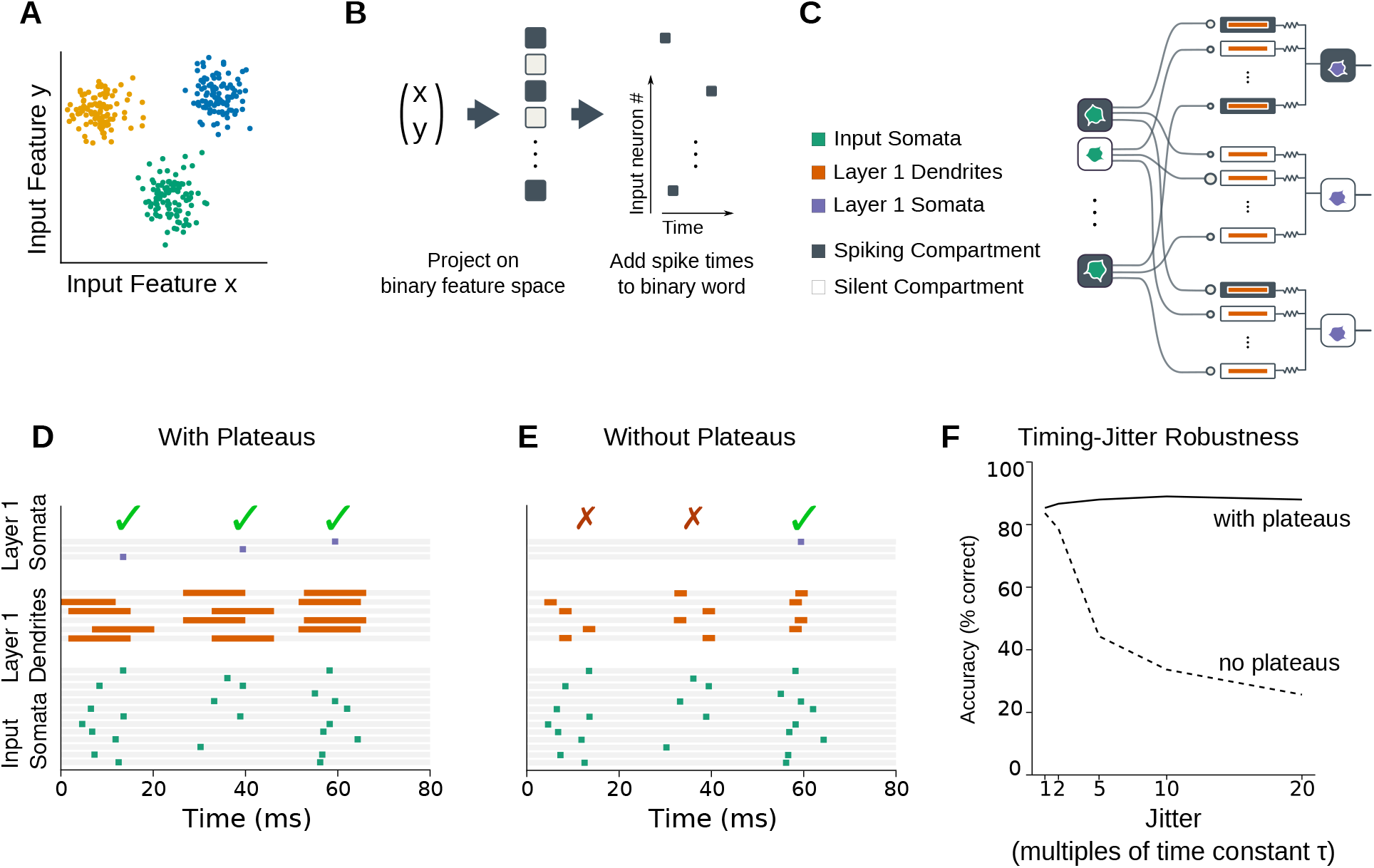
A simple conductance based model displays the same qualitative behavior as a detailed biophysical model. **(A)** The classification task performed by the spiking network of figures D, E, F. Each point is a 2D input vector *x*, the colors represent the different classes. **(B)** Procedure of transforming continuous 2D inputs *x* into input spikes for the network. First, *x* is projected onto a binary feature space to obtain a binary vector in a higher dimensional space. Then, spike times are added to this binary vector to produce the series of input spikes. For details, see subsection 3.4. **(C)** Schematic of the network architecture. The input somas spike according to the spike times obtained from the binary input vector. Each soma in the next layer has one dendrite per upstream soma, and each dendrite is connected to both one downstream and one upstream soma only. The dendrite-soma coupling is a bidirectional passive resistive coupling, whereas the upstream somas have a one-directional synaptic coupling onto the dendrites. **(D)** Example of the spiking network equipped with plateaus in the dendrites receiving asynchronous input spikes. It classifies the three inputs correctly in spite of the asynchrony. **(E)** Example of the spiking network equipped with dendrites without plateaus receiving asynchronous input spikes. The first two points are classified incorrectly, the network gets the third answer correct. **(F)** Summary of how well the network with plateaus and the network without plateaus deal with asynchrony *τ*, with the performance measured as the percentage of points of the classification task classified correctly. Without plateaus the performance drops off quickly (dashed line), whereas the network with plateaus does not suffer from performance degradation for this range of *τ* (solid line).

The network architecture is set up so that each dendrite is connected to both a unique upstream neuron and a unique downstream soma (see Figure 4C for a sketch). The assumption that each neuron connects to one dendrite of a downstream neuron is actually grounded in physiology (but see [7]), although it may appear like a strong assumption at a first glance: related inputs arrive at local clusters of spines synchronously [68]. We have modelled these dendritic patches that synchronously get excited by correlated inputs as one dendritic compartment. When a sufficiently large number of dendritic compartments have been excited, the soma will spike.

We have not explicitly accounted for inhibition in this model, although it plays an important role in determining when neurons fire. Our model tackles the question of how to deal with spike timing jitter in excitatory spikes. In order for it to be consistent, inhibition should at least not decrease spike timing synchrony. This is a reasonable assumption as inhibition plays an important role in improving synchronicity of spikes [15, 72]. Moreover, inhibition often follows excitatory inputs closely in time [31]. In our model, inhibition can be thought of as being implicitly accounted for, by regarding the inputs to the neurons as the net excitation in the presence of inhibition.

We transplanted the weight matrices of the trained BNN onto the spiking network, thereby obtaining a spiking network that could perform the classification. When the input neurons all spike simultaneously, the spiking network mimics the BNN exactly, i.e. the same units are active in both. But when asychrony is introduced, a discrepancy can arise. We introduce asynchrony in the network by moving the timing of the input spikes. Two examples of the spiking network receiving jittered input spikes are shown in Figure 4D and E. In Figure 4D the dendrites are furnished with plateau potentials (as in Figure 2F, top). These three input vectors are classified correctly despite the input spike jitter. In fact, the identity of the neurons spiking are still the same units that emit a 1 in its BNN counterpart.

This stands in contrast with the network performance without dendritic plateaus (Figure 4E), as in Figure 2F, bottom. In Figure 4E it can be seen that the network now fails to process two of the three input vectors correctly. The duration of the dendritic spikes is too short so that they are too far apart in time, and the soma fails to reach threshold. To test how quickly this degrades performance, we tested the accuracy (as percentage of inputs classified correctly) of the network as the asynchrony increased. The network with active dendrites coped well, but the performance of the network without dendrites with plateau potentials degraded rapidly. This is quantified in Figure 4F, where we see that classification accuracy drops precipitously if spike timing jitter exceeds the membrane time constant significantly. In contrast, dendritic plateaus maintain performance even when spike jitter exceeds membrane time constant by an order of magnitude. It is worth pointing out that spikes can occur in any unit as soon as threshold is reached; dendritic potentials do not therefore prevent single computations occurring more rapidly.

Together, these results show in principle how a cellular mechanism that captures the essential abstract features of dendritic spikes can serve to enhance robustness of non-trivial spiking computations in a network. Furthermore, they provide an abstract interpretation of rapid spiking computations as binary neural network computations.

## 2. Discussion

An animal’s survival can hinge on its ability to make rapid, reliable decisions. Consequently, there will be evolutionary pressure for neural circuits to function at the most rapid timescale possible in some situations. For example, studies of primate visual reaction time estimate each neuron in the synaptic pathway has approximately 10 ms to make a firing decision, a time window allowing 1–2 spikes in each unit on average [70, 69]. This places the excitatory units in these pathways in an effectively binary regime: either a neuron fires once during the entire computation or it does not.

We asked how cortical neurons might exploit dendritic nonlinearities to make such binary computations feasible in the biologically realistic situation of spike timing jitter and signalling noise. The significance of our work is to show that sparse yet reliable spiking computations may not require precisely synchronized inputs, yet yield and propagate reliable outputs.

Traditionally thought to be passive cables, we now know that dendrites possess a zoo of voltage-gated ion channels that generate nonlinear membrane potential dynamics [67, 61, 46, 63, 40, 50]. As with axonal conduction, dendritic excitability provides a means for signals to overcome spatial attenuation—so it is perhaps not surprising to find regenerative currents in dendrites, particularly the long, thin dendrites of cortical neurons.

It is far less obvious why dendritic action currents are so much slower than their axonal counterparts. Their temporal dynamics, along with their nonlinear amplitude dependence opens numerous ways for neurons to process time-varying signals. For instance, the dendrites of pyramidal neurons can perform complex tasks such as the discrimination of temporal signals [9] or the detection of coincident inputs [44]. In parallel with providing rich signal processing capabilities, dendritic transients influence activity-dependent synaptic plasticity [29, 39], and may allow neural circuits to learn temporal patterns [36, 35, 30].

We considered a complementary role for dendritic action currents that is not in conflict with any of these ideas, yet it addresses an outlying problem we believe is essential: making rapid cortical computation robust. Events in the external world are often asynchronous, yet may need to be processed by neural circuits that lump decisions into well-defined intervals. Inside the brain, conduction delays and intrinsic noise make asynchrony unavoidable in communication between neural circuits [19]. This poses a fundamental problem for the integration of related inputs: neurons with short membrane time constants can only integrate coincident inputs that arrive simultaneously within *∼* 1 ms of one another. Coordination of many such inputs in a distributed network therefore presents a fundamental challenge that was recognised in early attempts to understand computation in the brain [73]. We have shown that the long duration of nonlinear dendritic transients can remedy this situation by widening the integration time window of neurons.

Our hypothesis is consistent with several experimental findings. Blocking NMDA receptors impairs precise spike-timing synchrony and inter-area communication [75], indicating an important role for NMDA-dependent dendritic transients in facilitating coordinated communication between neural circuits. Recently it was shown that NMDA spikes are a major determinant of neuronal output in vivo, and that these dendritic spikes can be triggered by a handful of synaptic inputs [26, 51, 64, 11]. Our work provides further experimentally testable hypotheses: finer manipulation of dendritic transient time course, for example, by expression of chimaeric NMDA receptors [55] or calcium buffers, can shorten the integration window in cortical circuits. This should result in propagation failures, and failure to integrate distributed information. These effects should be visible directly in neural population activity and in behavioural performance in tasks involving rapid sensory discrimination, or fusion of information over multiple brain areas. Such an experiment would need to titrate the perturbation of dendritic dynamics to avoid less subtle circuit level effects that are seen in complete pharmacological blockade. This kind of manipulation would be exciting to apply in less extensively studied species, such as turtle, where cortical dynamics seem to operate close to a synfire chain regime [32], propagating distributed patterns in which participating neurons fire sparsely.

We have been careful to respect the essence of basic physiological facts when constructing our abstract model, but a tractable model often involves approximations. One conspicuous omission is to account for temporal variation in inhibition, which plays an important role in determining when and if spikes can fire in a network. Leaving aside simplicity, we have two justifications for this in the present work: first, we wanted to isolate a mechanistic ‘kernel’ for dealing with spiking jitter in excitatory input by assuming that inhibition is present, and, at minimum, not making matters worse. In this setting one may interpret the excitatory inputs to the abstract model in the network (Figure 4) as a *net* drive in the presence of both inhibition and excitation. We believe this is reasonable because inhibitory signals in many local circuits reflect local population activity and often reliably track excitatory input [31]. Secondly, overwhelming evidence shows that inhibition itself plays an important role in enhancing synchrony in neural populations [15, 72]. We want to return to the question of integrating these features of the physiology in future work, our hypothesis being that integrate-and-hold can serve to improve computational robustness, as we have shown, and furthermore permit information to be preserved throughout the phase of prominent network level oscillations in the brain that are largely orchestrated by inhibition.

Our model includes dynamics that are often simplified away in computational and theoretical studies. Much effort has been devoted to finding out the extent to which neuronal noise can deteriorate the function of neural networks [6, 22, 76]. Sometimes out of practical necessity, these studies may assume a point model of a neuron [24]. Our work shows that dendritic compartments and their dynamics may reduce the apparent fragility of spiking computations, prompting caution in how we interpret modelling results that lack these dynamics.

Other, simpler solutions to the asynchrony problem have been proposed. On possibility is that neural circuits exploit population averaging to overcome spike timing jitter [21, 65, 60, 43, 14]. However, this this is energetically costly [5] and would amount to scaling up the number of cells in a network to perform computations that could, in principle, be performed by fewer (but more robust) single units. We have argued that NMDA currents in distal dendrites can achieve robust, reliable threshold-based computation using relatively few resources. The longer duration of these potentials confers robustness to input timing without additional learning, reducing the number of neurons that must spike to achieve reliable signal transmission.

Synchrony could potentially be maintained in networks that are organized as feed-forward “synfire chains”, with relatively homogeneous transmission delays between nodes in each “rung” [1, 33]. [41] emphasize a role for refractoriness in maintaining synchrony, noting that post-spike inhibition “clips” late inputs, thereby maintaining a localized packet in time. [37] explore further the importance of dendritic nonlinearities in stabilizing packet synchrony. The extent to which spike timing is reliable remains an open question, one that likely depends on the circuit and context. An alternative approach to reliable spiking computations is for units to respond to specific, predictable, patterns of asynchronous inputs. Several computational studies have shown this is possible [13, 28, 53, 30]. However, these solutions are by design sensitive, rather than invariant, to the timing and order of inputs. Such solutions are thus less suited to the setting we considered here. We suggest that the contrasting properties of timing-sensitive and timing-robust architectures are likely both present in nervous systems, depending on the specific function of the circuits in question.

Fast action potentials allow rapid, massively parallel communication in the brain. In principle this gives spiking networks the ability to perform complex computations efficiently. However, decades of research has identified obstacles to implementing spiking computations under biologically realistic conditions. Interestingly, advances in neuromorphic hardware have revitalized attention for the question of how biology solves the problem. Our work shows a means for rapid spiking computations to function robustly by providing a resettable temporal buffer in the input to each spiking unit. We would be excited to see experimental tests of whether dendrites do indeed fulfil this function in the nervous system, and whether this simple principle offers a bio-inspired means to scale up reliable spiking computations in hardware implementations of spiking neural networks.

## 3. Methods

The code for running the simulations can be found in this GitHub repository:https://github.com/olearylab/active_dendrites_robust_spiking.

### 3.1 Biophysical model

The biophysical model used to test the plausibility of the abstract model was taken from [23], and was implemented in NEURON (version 7.5) [34]. The model can be found on ModelDB (no. 249705). The model represented a layer 5 pyramidal neuron [2], and consisted of 85 compartments and 439 segments; 36 compartments for basal dendrites, 45 for apical dendrites, 3 for the soma, and one for the axon. A range of voltage-gated ion channels is included: sodium, A-type potassium, both high-voltage and low-voltage gated calcium, HCN, calcium-activated potassium, and K_v_ type channels are present.

The glutamate release is simulated at basal dendrites with the indices 14, 15, and 34. These indices were chosen for no particular reason; any collection of numbers would have worked. The glutamate stimuli were all given a fraction of 0.9 away from the soma (where 1 is the total length of the dendrite), and were separated in time from each other with a 50ms delay.

For Figure 2G, a passive inhibitory current was added to basal dendrite 34 in which an NMDA spike was triggered. The reversal potential of this current was that of GABA_A_, i.e. -80 mV. The maximal conductance of this current was simulated as 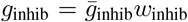, with 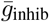 being the maximal conductance of 0.001 mS / cm^2^, and *w*_inhib_ the weight that was varied during the simulation, such that *w* = {0.0, 1.0, 2.5, 2.0, 3.0, 3.5, 4.0}. In Figure 2G we plotted against the dimensionless quantity *g*_inhib_*/g*_NMDA_, where *g*_NMDA_ is 0.005 mS / cm^2^.

### 3.2 Abstract model

The simplified model in Figure 2E,F is described by two differential equations for each dendritic branch, and two for the soma. The dynamics of the dendritic membrane potential *V* ^*d*^ and somatic potential *V* ^*s*^ are given by

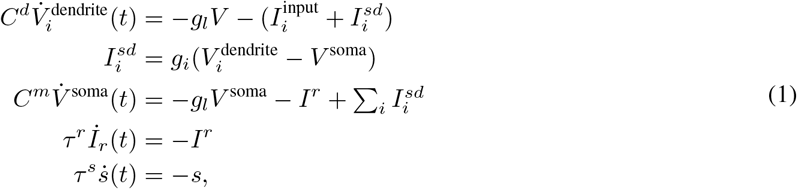

where *g*_*l*_ is the leak conductance, 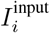 is the current triggered by a spike arriving at dendrite *i*, 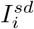 is the current flowing between dendrite *i* and the soma, with *g*_*i*_ the conductance between the two, *τ*^*x*^ is the time constant of variable *x, I*^*r*^ is a refractory current, and *s* is the postsynaptic conductance of the neuron. When the dendrite reaches threshold Θ^dendrite^ the dendrite remains at threshold for *P* ms:

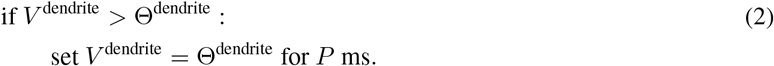

A spike is triggered when the soma reaches its threshold Θ^soma^:

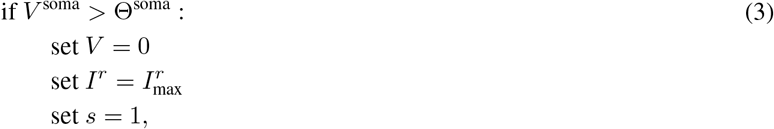

i.e. the membrane potential is reset, a refractory current is activated, and the postsynaptic conductance increases.

Unless mentioned otherwise, the parameter values that were used in simulations are given in Table 1.

**Table 1:** Standard values of parameters used.

| $C^d$ | $C^m$ | $\tau^r$ | $\tau^s$ | $g_l$ | $g_i$ | $\Theta^d$ | $\Theta^s$ | $I_{\max}^r$ | $V_E$ |
| --- | --- | --- | --- | --- | --- | --- | --- | --- | --- |
| 1 nF | 0.5 nF | 10 ms | 10 ms | $0.05 \mu\text{S}$ | $1 \mu\text{S}$ | 1 mV | 1 mV | 5 nA | 100mV |

For Figure 2E,F, for the case with active dendrites, we used this abstract model, furnished with three dendritic compartments. The dendrites were given boxcar functions, with an amplitude of 50 and a width of 1 ms, as input pulses:

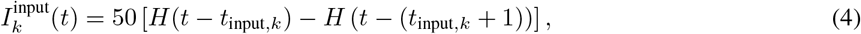

and *k* ∈ {1, 2, 3}. These input pulses were spaced 10 ms apart; *t*_input,*i*_ = 10*k* ms. Each pulse is strong enough by itself to trigger the plateau in each dendrite, thereby extending its duration. Taken together these three plateaus are strong enough to trigger a spike in the soma. For Figure 2E,F, for the case without dendrites, these pulses were fed to the soma directly. Due to the lack of plateaus they do not sum and fail to trigger a spike.

### 3.3 Neuron with asynchronous inputs

For Figure 3 the abstract model was used. The spike times of the incoming spikes were drawn from a normal distribution. The jitter of the spike times was defined as the standard deviation of this distribution. This jitter should always be compared with the membrane time constant *τ* of the neuron, for which we used 10 ms.

We used jitter values of 1*τ* for the synchronous case and 10*τ* for the asynchronous case respectively, i.e. *t*_input_ *∼ 𝒩* (0, *ατ*), *α* ∈ {10, 100}.

For all the computed values 500 simulations with a single neuron and randomly initialized spike times were performed. During each simulation, the neuron received 10 input spikes. Voltage trajectories were normalized with respect to the mean voltage; that is, the mean *µ*(*t*) was computed, and the normalized voltage trajectories were then given by 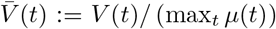. For Figure 3B, the solid line is the mean voltage over the ensemble of simulations, and the shaded region is the inner 90% of the voltage distribution.

In Figure 3C,D,E all variables were computed inside the window of spikes arriving. The peak depolarization plotted is the mean of the ensemble of peak depolarizations for each simulation. Similarly, the coefficient of variation *C*_*v*_ plotted is the *C*_*v*_ for each jitter value, where that particular *C*_*v*_ is computed over all 500 simulations for that jitter value.

For Figure 3E the minimum number of input spikes was computed that would push *C*_*v*_ down to 0.1 for a particular value of the jitter value.

### 3.4 Binary network

The data consisted of 2D points which were assigned to different classes. The three classes were Gaussian clusters, with means *µ*_*i*_ = {(4, 5), (*−*12, 2), (10, *−*7)} respectively, and isotropic covariance of *σ* = 1.5. 100 points *x*_*i*_ per class *i* were generated from *x*_*i*_ *∼ 𝒩* (*µ*_*i*_, *σ*).

The task of the binary neural network (BNN) was to classify the points correctly. Because we wanted to interpret the continuous 2D points as input spikes to our network, we binarized the data first. To this end the input vectors *x* ∈ ℝ^2^ were transformed by mapping them onto {0, 1}^13^, i.e. every point was mapped onto a 13 dimensional vector of ones and zeros. This was achieved by randomly generating *W ∼ 𝒩*_13_(**0**, *I*) and *b ∼ 𝒩* (0, 1). Then we applied

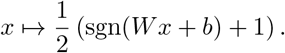

To train our binary network we made use of surrogate gradients [53]. In short, we defined a function *ϕ* that for the forward pass (i.e. the network unit outputs) acted like a step function

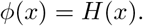

But for the purposes of backpropagation the derivative of *ϕ* is defined to be

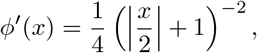

which is the superspike derivative [74], equivalent to the derivative of a fast sigmoid. Using this derivative allows us to train the network, in spite of the step function having a derivative that is zero almost everywhere.

The network weights were initialized with the distribution:

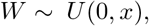

where 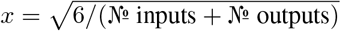.

The biases of the BNN units were set to *−*1, to enforce a positive firing threshold for the neurons.

Each dendrite receives input from one presynaptic neuron. Let *N*_*d*_ be the number of dendrites per neuron for a layer, and *N*_*o*_ the number of somas. First an *N*_*d*_ *× N*_*d*_ diagonal matrix is constructed for each soma in the next layer. Then the total weight matrix will be the vertical concatenation of all these diagonal matrices.

Similarly, each dendrite is coupled with only one soma. This is achieved by constructing an *N*_*o*_ *× N*_*d*_ matrix for output unit *i* where only the *i*th row is non-zero. The overall weight matrix is the horizontal concatenation of these matrices.

The weight matrices used by the network alternate between the matrices with ‘dendrites constraints’ (a matrix that projects onto a layer of dendrites), and matrices with ‘soma constraints’ (a matrix projecting onto a layer of somas).

For both weight matrices elements that are initially zero, remain zero during the training procedure, and the elements of the weight matrices were constrained to be nonnegative, by having the activation of a unit in the BNN be

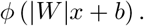

We implemented this constraint because we modelled pyramidal neurons, which are excitatory neurons. Therefore, the weights from the connections of these neurons should be positive.

The BNN was trained using stochastic gradient descent and surrogate gradient methods. First the network outputs *ŷ*(*x*) where put through the softmax function

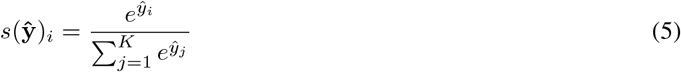

which was then parsed through the cross-entropy function

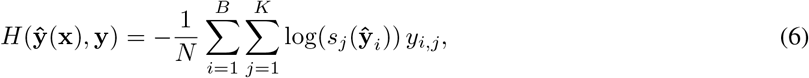

where ŷ_*i*_ is the output in response to input x_*i*_ belonging to the batch of size *B* and *y*_*i,j*_ the *j*th element of the corresponding target.

The stochastic gradient descent was performed by randomly shuffling the inputs and labels and choosing a batch size (30 in our case). Then in each epoch all batches were iterated over, and the parameters were updated using the Adam algorithm. Training was terminated when the network gave the correct answer for more than 90% of all input points.

### 3.5 Spiking network

After having a set of weights that gave good performance with the BNN those weights were transplanted to a spiking network with the abstract models from subsection 3.2. This was done through setting the input currents to

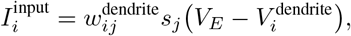

where *s*_*j*_(*t*) is the postsynaptic conductance of neuron *j*, which projects on dendrites *i* with weight 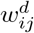. Here, 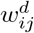 is the nonzero element of the *i*th row of the weight matrix of this layer, trained with the ‘dendrite constraints’ (see subsection 3.4). This element is unique due to the constraints imposed on the weight matrices during training.

This dendrite is coupled to its soma with weight 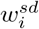 to give the current

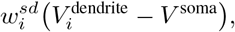

with 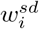 the (again unique) nonzero element of the *i*th row of the next weight matrix, trained with the ‘soma constraints’, and so on.

The 13 dimensional spikes were fed into the network by means of an input layer. This layer consisted of 13 neurons, one for each element in the binary input vectors. If the corresponding element was a 1, this neuron would be made to fire by giving them the same boxcar-shaped input current as described in subsection 3.2.

The degree of synchrony in the input spikes could then be controlled by varying the timing at which these impulses were given to the input neurons. Every input spike packet was centered around an input time 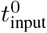. Then the degree of asynchrony *τ* was varied by letting each spiketime, for each input neuron *i*, be

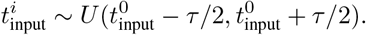

To compare the performance of a spiking network without the dendritic plateaus to networks with plateaus, the ‘hold’ on the dynamics of 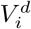 was removed. Therefore, the dendrite potential would decay immediately back to rest after reaching threshold. Otherwise, the architecture remained unchanged. For Figure 4D,E the input volleys were centered at 15ms, 35ms, and 55ms, with an asynchrony measure of *τ* = 10ms.

An answer given by the spiking network was considered correct if neuron *i* in the output layer produced one spike when the input point belonged to class *i*, and the other neurons remained silent.

To compare the accuracies with and without plateaus in Figure 4F, the spike times of the input neurons were produced independently for each data point and each value of *τ*, with the spike packet being centered at 10 ms. Then the same spike times were used for both network versions, and the accuracy of the network was measured as the percentage of points classified correctly by the network. In each simulation, each network saw one input vector only, to prevent interference from multiple overlapping inputs for high jitter values.

## 4. Acknowledgements

This work was supported by ERC grant 716643 FLEXNEURO (TO), an Engineering and Physical Sciences Research Council DTP studentship (TJSB), and Isaac Newton Trust and Leverhulme Fellowship ECF-2020-352 (MER).

